# HLA-DQ2/8 genotype is associated with strain-level gut microbiome divergence and lower serum pantothenate in healthy adults

**DOI:** 10.1101/2025.10.20.683361

**Authors:** Chuanwen Kan, Mo Hu, Ning Wang, Qianhui Zhang, Lingxiao Chaihu, Xuan Jiang, Chun Wang, Wenwei Lu, Guanbo Wang, Mingkun Li, Li Zhang

## Abstract

HLA-DQ2/8 haplotypes are established genetic risk factors for autoimmune diseases and are known to influence gut microbiota assembly in early life. However, their impact on the adult microbiome and functional consequences for host physiology remain unclear. Here, we performed a genotype-stratified multi-omics analysis of 60 healthy adults, including 28 HLA-DQ2/8 carriers and 32 non-carriers. We found that host HLA-DQ2/8 genotype was significantly associated with gut microbiome composition, with an effect size exceeding that of sex and BMI. HLA-DQ2/8 carriers exhibited higher gut microbial alpha-diversity, lower virulence factor abundance, and a distinct species profile enriched in butyrate-producing taxa. We identified pervasive intra-species phylogenetic and functional divergence linked to the DQ2/8 genotype. This diversification reflects predicted HLA-restricted microbial peptide-binding specificities, suggesting a possible role for antigen presentation-mediated immune selection, a mechanism further supported by AlphaFold3 structural modeling. We found an enrichment of microbial pathways for pantothenate and coenzyme A (CoA) biosynthesis in carriers, primarily driven by functionally divergent *Blautia obeum* strains. This functional shift paralleled lower levels of serum pantothenate and HDL-cholesterol in the host. Our findings suggest a potential genotype–microbiome–host axis where antigen presentation-mediated immune selection may modulate microbial adaptation, with possible implications for the host availability of essential cofactor precursors and lipid metabolism.

## Introduction

Autoimmune diseases such as celiac disease, type 1 diabetes, and autoimmune thyroid disorders are strongly associated with both HLA-DQ genotypes and gut microbiome alterations ^[1–4]^. The HLA-DQ genes (HLA-DQA1 and HLA-DQB1) encode major histocompatibility complex (MHC) class II heterodimers expressed on antigen-presenting cells. These proteins present dietary and microbial peptides to CD4⁺ T cells, thereby orchestrating adaptive immune responses. Among these, the DQ2.5, DQ2.2, and DQ8 haplotypes confer the highest genetic risk for celiac disease, being present in >95% of cases, and also increase susceptibility to multiple autoimmune diseases. However, the low penetrance of disease among carriers underscores that additional environmental and microbial factors modulate risk ^[5,6]^.

Accumulating evidence indicates that HLA polymorphisms influence human gut microbial communities even in asymptomatic individuals. HLA variation has been linked to microbiome community structure across body sites ^[7]^, and the HUNT microbiome GWAS recently identified and independently replicated an association between rs28407950 at the HLA-DQB1 locus and *Agathobacter* sp000434275 abundance ^[8]^. Rodent models expressing human HLA alleles further support a causal influence on the gut microbiome and on autoimmune phenotypes ^[9,10]^. Focusing on HLA-DQ2/8, human cohort studies first reported microbiome signatures in high-risk infants, where DQ2 carriage was associated with reduced *Bifidobacterium* and disrupted co-occurrence networks by 4 months, before gluten exposure ^[3]^. These findings were subsequently extended to the general population in the ABIS cohort, where genotype-associated microbiome profiles were evident at age one ^[11]^, and microbial markers predictive of future celiac disease were identified at that same age ^[4]^. Later studies confirmed similar patterns in schoolchildren, among whom DQ2/8 carriers exhibited reduced phylogenetic diversity with distinct genus-level signatures ^[12]^. More recently, functional metagenomics has shown that DQ2/8 genotypes were linked to the enrichment of microbial pathways with inflammatory potential, indicating that HLA–microbiome interactions extend beyond taxonomy to functional capacity ^[13]^. Together, these findings highlight a role for HLA-DQ2/8 in modulating the gut microbiome from early life, with potential consequences for immune development and autoimmune risk trajectories.

Despite these advances, important gaps remain. Investigations to date have focused on infants and children, leaving the impact of HLA-DQ2/8 genotype on the adult microbiome largely unexplored, even though these genotypes persist throughout life and the microbiome remains dynamic across the lifespan. Furthermore, most prior studies have concentrated on taxonomic composition, neglecting functional pathways, strain-level phylogenetic divergence, or connections to host metabolism and phenotypes. These dimensions are particularly relevant to autoimmunity because microbial strain variation and functional capacity may determine the antigenic repertoire, inflammatory potential, and metabolite output encountered by the host immune system. Few studies have adequately controlled for confounders such as diet, antibiotics, and lifestyle, which strongly influence the microbiome. As a result, the independent contributions of HLA-DQ2/8 genotype to microbial ecology and host–microbe interactions remain poorly defined.

To address these gaps, we conducted a multi-omics study of 60 healthy young adults, comparing HLA-DQ2/8 carriers and non-carriers. Our design integrated shotgun metagenomic sequencing of stool and saliva, untargeted serum metabolomics, clinical phenotyping, questionnaire, and detailed dietary nutrient intake assessments, enabling high-resolution analyses while accounting for lifestyle and environmental covariates. By linking HLA-DQ2/8 genotypes to microbial functions, strain-level diversity, and host physiology, our study provides new insights into how host genetics may influence the functional landscape of the gut microbiome in health.

## Results

### Participant Stratification and Assessment of Cohort Homogeneity

This investigation represents a genotype-stratified, observational analysis nested within a parent single-arm, ABA dietary intervention cohort. A total of 60 healthy adults were genotyped for HLA-DQ. The cohort included carriers of single haplotypes DQ2.2 (n=14), DQ2.5 (n=8), and DQ8 (n=5), as well as one individual with combined genotype DQ2.2/8. Based on this distribution, participants were stratified into 28 HLA-DQ2/8 carriers (DQ2/8+) and 32 non-carriers (DQ2/8–). All participants were negative for tTG IgA, ruling out active celiac-related autoimmunity.

To achieve a robust characterization of individual features while accounting for the parent study’s dietary phases, we utilized per-person averages calculated across the five sampling timepoints. This approach effectively minimized intra-individual noise and transient fluctuations, providing a sustained, high-resolution representation of the host–microbiome landscape for genotype-based comparisons.

The homogeneity of the two groups was rigorously confirmed using these aggregated measures. No significant differences were observed between DQ2/8+ and DQ2/8–groups in primary demographics, including age, sex, BMI, and ethnicity (Table 1). Notably, the groups were well-matched in their habitual dietary patterns, encompassing mean daily intake of total calories, macronutrients, fiber, micronutrients, and dietary diversity (Table 1). Further evaluation of physiological, biochemical, and psychosocial parameters, including body composition, liver/renal function, serum proteins, red blood cell indices, and self-reported wellness indicators, confirmed the lack of disparities between the two groups (Supplementary Table 1). These results establish a well-balanced cohort, providing a rigorous baseline for investigating the downstream impact of HLA-DQ2/8 on the microbiome-host axis.

**Table 1.** Demographic characteristics, dietary diversity, and daily nutrient intake of the study population.

|  | DQ2/8– (n=32) | DQ2/8+ (n=28) | P value* |
| --- | --- | --- | --- |
| **Demographics** |  |  |  |
| Age (years) | 24.0 [22.8-25.0] | 24.5 [23.0-25.0] | 0.15 |
| Male sex | 4 (12.5%) | 6 (21.4%) | 0.36 |
| BMI | 20.9 [19.4-21.5] | 21.0 [19.9-22.3] | 0.32 |
| Ethnicity |  |  | 0.33 |
| Han | 31 (96.9%) | 26 (92.9%) |  |
| Other | 1 (3.1%) | 2 (7.1%) |  |
| **Dietary diversity** |  |  |  |
| Number of food groups | 7.7 [7.0-8.0] | 8.0 [7.2-8.4] | 0.284 |
| Number of food subgroups | 9.2 [8.6-9.7] | 9.8 [8.9-10.2] | 0.339 |
| **Daily nutrient intake** |  |  |  |
| Calorie (kcal) | 1324.9 [1182.9-1463.9] | 1287.8 [1142.1-1579.2] | 0.92 |
| Fat (g) | 55.8 [47.5-62.7] | 55.5 [45.7-63.1] | 0.68 |
| Protein (g) | 67.4 [60.2-78.2] | 68.4 [60.8-81.6] | 0.61 |
| Carbohydrate (g) | 134.5 [126.1-175.8] | 143.5 [127.4-160.6] | 0.53 |
| Fiber (g) | 6.0 [4.9-7.7] | 5.7 [5.0-6.8] | 0.68 |
| Cholesterol (mg) | 469.7 [392.4-614.3] | 487.2 [380.0-533.7] | 0.57 |
| Calcium (mg) | 284.1 [244.6-384.3] | 251.0 [220.9-308.4] | 0.16 |
| Sodium (mg) | 1865.2 [1703.7-2239.3] | 1916.5 [1809.7-2198.1] | 0.59 |
| Vitamin A (μg) | 183.7 [156.2-244.6] | 172.9 [139.4-219.4] | 0.19 |
| Vitamin B1 (mg) | 0.5 [0.5-0.6] | 0.5 [0.4-0.6] | 0.34 |
| Vitamin B2 (mg) | 0.7 [0.6-0.8] | 0.6 [0.5-0.8] | 0.16 |
| Vitamin B3 (mg) | 13.4 [11.5-14.6] | 13.8 [12.1-18.1] | 0.15 |
| Vitamin C (mg) | 53.5 [35.3-78.6] | 56.6 [40.5-70.1] | 0.69 |
| Vitamin E (mg) | 24.5 [21.1-28.9] | 24.7 [21.6-29.6] | 0.74 |

\*The p value for ethnicity was calculated using Fisher’s exact test, while p values for other factors were calculated using the Wilcoxon rank-sum test. Categorical variables are shown as number (percentage), and the rest data are presented as median [IQR].

### Gut microbial diversity, taxonomic composition, and functional pathways vary by HLA-DQ2/8 genotype

Gut microbial α-diversity (Shannon index) was significantly higher in DQ2/8+ individuals (Figure 1A). They also had a lower abundance of virulence genes (Figure 1B). PERMANOVA indicated that the HLA-DQ genotype was significantly associated with gut species composition (p<0.05, R^2^=0.03, Figure 1C; Figure S2). While dietary factors, particularly fiber intake, remained the dominant drivers of microbial variation, the estimated contribution of genotype exceeded that of sex and BMI (Figure 1C; Figure S2).

**Figure 1.**
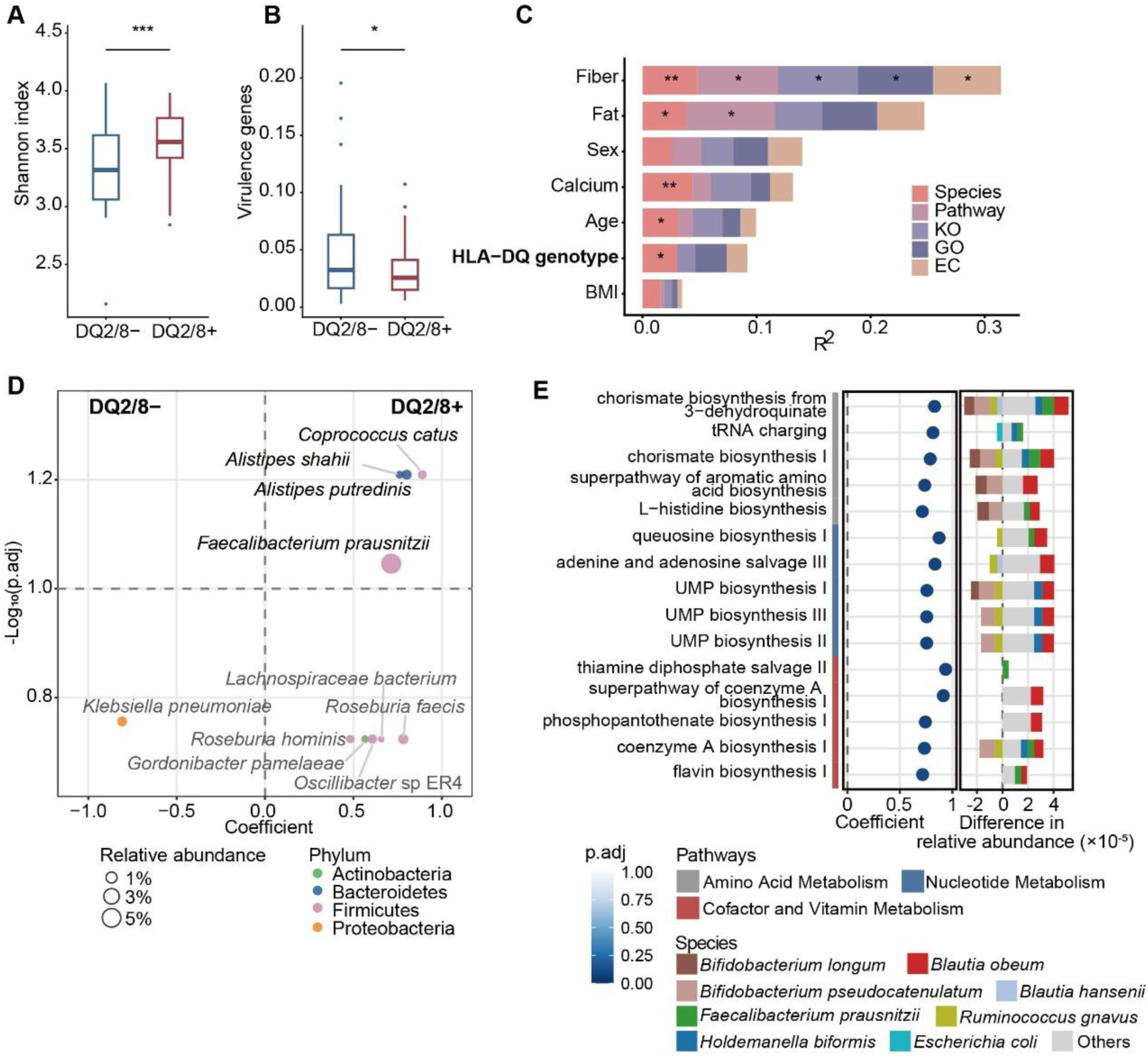
Gut microbial diversity, composition, and functional pathways vary by HLA-DQ2/8 genotype. (A–B) Comparison of (A) Shannon index and (B) bacterial virulence gene abundance between DQ2/8+ and DQ2/8– individuals. Boxplots show the median (center line), interquartile range (box), and 1.5×IQR (whiskers). (C) Covariates associated with gut microbiome variation, ranked by Adonis R² values across five microbiome profiles (PERMANOVA). Dietary features significantly associated with taxonomic profiles, as well as demographic characteristics are shown. For the full results encompassing all 12 analyzed features, see Figure S2. (D) Species differentially abundant between groups. (E) Microbial pathways differentially abundant between groups. Left: Differential pathways with p.adj < 0.1. Right: Top 8 contributing species. In (A–C), asterisks indicate significance (*p<0.05; **p<0.01; ***p<0.001). In (A, B, D, E), p values and effect estimates were derived from multiple linear regression models adjusted for age, sex, BMI, and dietary nutrient intake. Positive coefficients indicate higher abundance in DQ2/8+, whereas negative coefficients indicate higher abundance in DQ2/8–.

Differential abundance analysis identified an enrichment of *Faecalibacterium prausnitzii*, *Coprococcus catus*, and members of the *Alistipes* genus in the DQ2/8+ group (Figure 1D). The former two are butyrate producers, and are recognized for their roles in anti-inflammatory signaling and the reinforcement of intestinal barrier integrity ^[14,15]^. Additionally, a trend toward enrichment was observed for another two butyrate producers *Roseburia faecis* and *R. hominis* (Figure 1D). Conversely, DQ2/8–individuals showed a notable trend toward the enrichment of *Klebsiella pneumoniae*, a well-documented opportunistic pathogen. This taxonomic divergence functionally aligns with the lower overall abundance of virulence genes observed in the DQ2/8+ group, suggesting a gut environment with reduced pathogenic potential.

Although global microbial functional profiles (pathways, KOs, GOs, ECs) did not differ significantly between the two groups (Figure 1C), specific pathways were genotype-associated (Figure 1E). DQ2/8+ carriers had an increased abundance of vitamin-related pathways, including thiamine diphosphate salvage (B1), flavin biosynthesis (B2), phosphopantothenate biosynthesis (B5), coenzyme A biosynthesis, and the superpathway of coenzyme A biosynthesis, largely contributed by *Blautia obeum*. Intriguingly, the abundance of *B. obeum* itself did not differ between groups. This discrepancy indicates that the observed functional enrichment is likely contributed by strain-level genomic variation.

In contrast to the gut, the salivary microbiome exhibited no difference in diversity, taxonomic composition, or functional potential between the DQ2/8+ and DQ2/8–individuals.

### HLA-DQ2/8 genotype is associated with intra-species phylogenetic divergence and functional specialization

To assess intra-species divergence, we applied phylogenetic generalized linear mixed models (pGLMM) in anpan ^[16,17]^. Significant genotype-linked phylogenetic divergence was detected in 15 gut species, 12 of which were Firmicutes (Figure 2A). In saliva, 10 divergent species were identified, predominantly *Streptococcus* (Figure S3). These findings indicate that HLA-DQ genotype may impose selective pressures on intra-species evolution.

**Figure 2.**
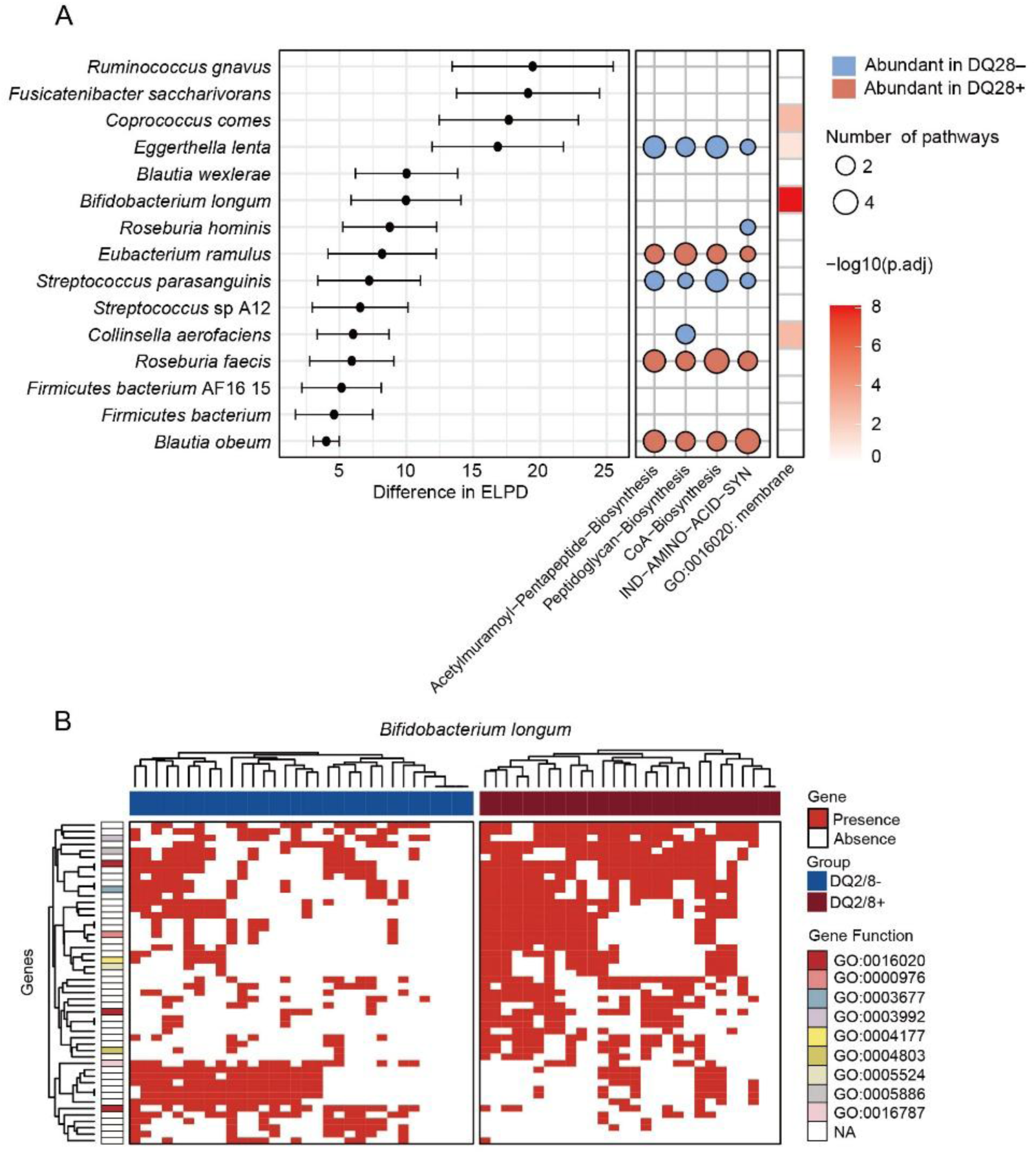
Intra-species phylogenetic divergence and functional specialization associated with HLA-DQ2/8 genotype. (A) Gut microbial intra-species divergence linked to DQ2/8 genotype. Left: Phylogenetic divergence between the DQ2/8+ and DQ2/8– groups, as indicated by differences in expected log pointwise predictive density (ΔELPD) computed by anpan. Species with ΔELPD > 4 are shown. Center: Pathways showing genotype-associated divergences across species. Only pathways detected in more than five significant pathway-species associations are displayed. Right: Heatmap of Gene Ontology (GO) enrichment of microbial genes showing differential prevalence between genotypes within individual taxa. Only GO terms with p.adj < 0.1 in at least two species are shown. (B) The top 50 *Bifidobacterium longum* gene families exhibiting the greatest prevalence differences between the genotypes. Each row represents a UniRef90 gene family and each column a sample. Genes are grouped by functional category, with membrane-related genes (GO 0016020) highlighted in red labels.

We next tested whether the observed phylogenetic divergence also manifested as functional differences, using pathway random effects models that controlled for species abundance in anpan. In the gut microbiome, four pathways showed the most significant genotype-associated divergence across multiple species, including *B. obeum* (Figure 2A): (i) the peptidoglycan biosynthesis pathway, which produces peptidoglycan as the predominant cell wall antigen of Gram-positive bacteria; (ii) the acetylmuramoyl-pentapeptide biosynthesis pathway, a critical precursor step for peptidoglycan formation; (iii) the IND-AMINO-ACID-SYN pathway, responsible for the synthesis of indole derivatives; and (iv) the CoA biosynthesis pathway. Notably, the CoA biosynthesis pathway had already emerged as differential in the overall gut microbiome analysis, with *B. obeum* contributing the largest share of pathway abundance (Figure 1E). This convergence highlights *B. obeum* as a principal contributor of CoA-related functional divergence between DQ2/8 carriers and non-carriers. In the salivary microbiome, genotype-associated shifts were observed in arginine biosynthesis, lactate fermentation, glycolysis variants, and de novo purine nucleotide biosynthesis within *S. salivarius* and *S. pneumoniae* (Figure S3).

To further investigate intra-species microbial adaptation to HLA-DQ genotype at a gene-level resolution, we identified 11,861 genes showing differential prevalence between the DQ2/8+ and DQ2/8– groups within individual taxa, followed by GO enrichment analysis. Remarkably, both gut and salivary microbiomes showed prominent enrichment of membrane-associated processes (GO:0016020, Figure 2A, Figure S3). This category includes genes involved in membrane protein folding, assembly, secretion, and transport. Together, these processes highlight adaptations related to the modification of surface epitopes and secreted factors that can be recognized by the host immune system, consistent with genotype-associated differences observed in microbial antigen biosynthesis pathways.

To provide a high-resolution visual demonstration of these genomic shifts, we focused on *Bifidobacterium longum*, the most prevalent gut species exhibiting significant phylogenetic divergence. By prioritizing the top 50 gene families with the most pronounced prevalence differences, we revealed a genotype-specific presence/absence pattern (Figure 2B). This case study presents a clear genomic signature of the segregating selective pressures exerted by the host HLA-DQ landscape at the strain level.

### Predicted HLA-DQ–mediated antigen presentation aligns with strain-level microbial divergence

To assess whether the intra-species distribution of microbial genes varied according to HLA-DQ genotype, we analyzed the predicted binding affinities of strain-specific membrane-associated gene products against six representative HLA-DQ molecules (DQ2.2, DQ2.5, DQ8, DQ6, DQ7.5, and DQ9). In the gut microbiome, we observed an inverse relationship: genes with strong or weak predicted binding were less prevalent among carriers of the corresponding HLA-DQ haplotype compared to non-binding genes (Figure 3A). This pattern indicates that HLA-DQ–restricted antigen presentation may effectively filter microbial strains based on their membrane-associated gene content. This relationship was absent in the salivary microbiome, suggesting a less prominent immune-microbe interface.

**Figure 3.**
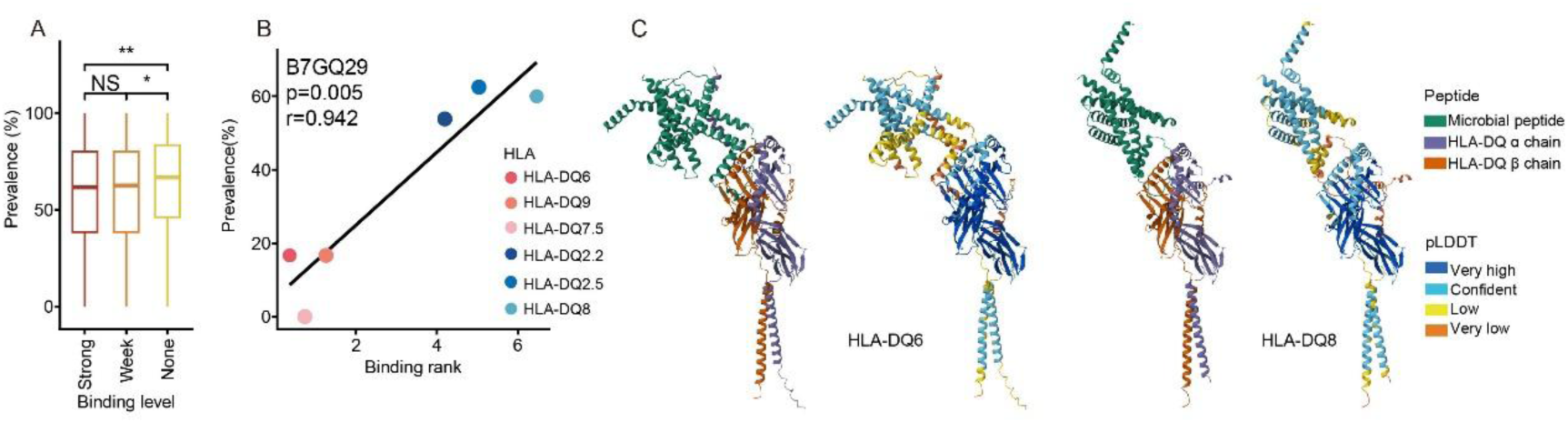
HLA-DQ binding affinity is associated with gut microbial gene prevalence. (A) Prevalence of microbial genes with different HLA-DQ binding affinities. The x-axis shows the predicted binding affinity of each gene product against six representative HLA-DQ heterodimers (DQ2.2, DQ2.5, DQ8, DQ6, DQ7.5, and DQ9). The binding affinity was predicted using NetMHCIIpan v4.3 and classified by the recommended %Rank thresholds as strong (≤ 1%), weak (1–5%), or non-binding (>5%). The y-axis shows the prevalence of each gene among isolated haplotype carriers, defined as individuals carrying the respective HLA-DQ haplotype without additional risk haplotypes. For DQ6 and DQ9, individuals were combined into a single reference group for prevalence calculation. A total of 608 differential genes encoding membrane proteins were analyzed. Boxplots show the median (center line), interquartile range (box), and 1.5×IQR (whiskers). Asterisks denote statistical significance from Wilcoxon tests (*p<0.05; **p<0.01). (B) The relationship between predicted binding affinity rank (lower=stronger) and gene prevalence among isolated haplotype carriers for a representative gene product, B7GQ29 from gut *B. longum*. P values were calculated using Spearman correlation. (C) AlphaFold3 structural models of microbial peptide–HLA-DQ interactions, illustrating stable binding of B7GQ29 to DQ6 versus weak binding to DQ8. Left: Peptide–HLA complex structures. Right: pLDDT confidence scores.

We further performed structural validation for high-confidence candidates exhibiting strong linear affinity–prevalence correlations. For instance, the gene product B7GQ29 from *B. longum* exhibited the highest predicted binding affinity to HLA-DQ6, the genotype where it was the rarest, and the weakest binding to DQ8, where it was most prevalent (Figure 3B). Structural modeling via AlphaFold3 ^[18]^ confirmed that B7GQ29 forms a stable complex within the DQ6 binding groove, whereas its interaction with DQ8 is characterized by poor complex stability (Figure 3C).

Together, these analyses support an association between HLA-DQ–restricted antigen presentation and microbial gene prevalence, consistent with a potential role for antigen presentation in modulating strain-level microbial adaptation to host genotype.

### HLA-DQ2/8 carriers show lower serum pantothenate and HDL-C

To determine whether genotype-associated microbial shifts were accompanied by systemic host alterations, we integrated our functional metagenomic data with serum metabolomics. Among the 59 metabolites theoretically linked to the differentially abundant microbial pathways (Supplementary Table 2), 14 were detected in the serum. Pantothenate emerged as the sole metabolite showing significant inter-group differences, with lower levels observed in DQ2/8+ carriers (Figure 4A).

**Figure 4.**
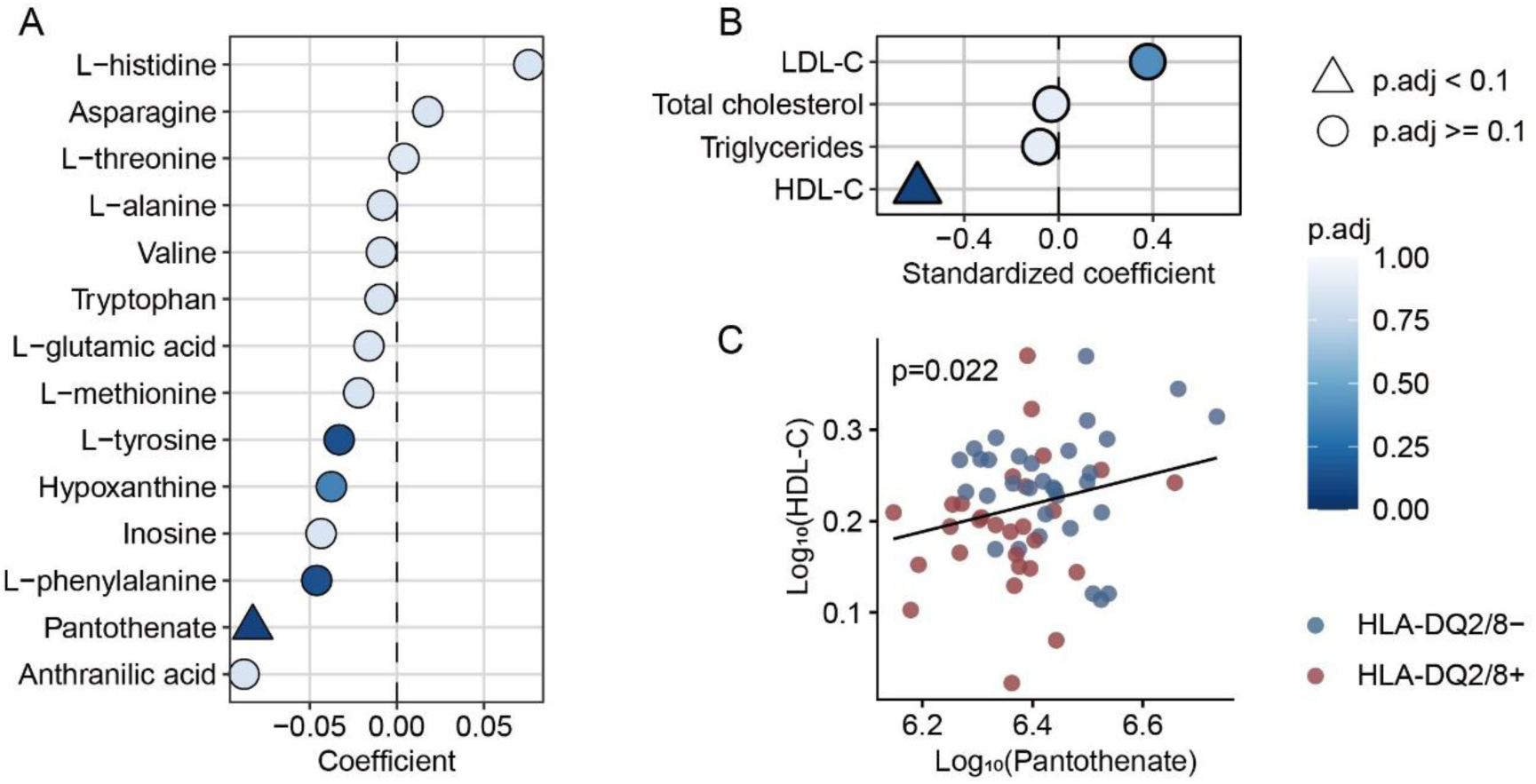
Host metabolic alterations associated with HLA-DQ2/8 genotype. (A) Serum metabolites mapped to differentially abundant microbial pathways. (B) Lipid-related phenotypes. (C) Correlations between serum pantothenate and HDL-C. Coefficient estimates and p values were derived from multiple linear regression models adjusted for age, sex and BMI. Positive coefficients indicate higher values in DQ2/8+, whereas negative coefficients indicate higher values in DQ2/8–.

Given that pantothenate is the obligate precursor of coenzyme A and a central regulator of lipid metabolism ^[19,20]^, we further evaluated host lipid-related clinical profiles. Congruent with the observed metabolite depletion, DQ2/8+ individuals exhibited significantly lower levels of high-density lipoprotein cholesterol (HDL-C) (Figure 4B). Correlation analysis confirmed a significant positive association between serum pantothenate concentrations and HDL-C levels (Figure 4C), suggesting a coordinated metabolic association.

Beyond lipid metabolism, no significant differences were detected in other phenotypic domains, including gut health markers, fecal short-chain fatty acids, cardiovascular parameters, blood glucose, immune cell profiles, or platelet indices (Supplementary Table 3).

### Longitudinal Stability and Robustness of Genotype-Associated Signatures

Given the longitudinal design of the dietary intervention, we performed a robustness check to ensure that the observed genotype associations were stable and not confounded by short-term dietary shifts. Linear mixed-effects models (LMM), incorporating the specific intervention phases (baseline, low-gluten, and high-gluten time points), revealed no significant genotype-by-diet interactions for the primary differentiating features across all omics layers. Specifically, the stability was confirmed for the gut microbial Shannon index, virulence genes, microbial species, pathways, serum pantothenate and HDL-C (Supplementary Table 4). These longitudinal validations demonstrate that the HLA-DQ2/8-associated effects remain invariant across varied dietary contexts, thereby robustly justifying our analytical approach of utilizing time-point averages for the primary cross-sectional comparisons.

## Discussion

Based on the differences in microbiomes, host phenotypes, and host-microbe interactions between healthy HLA-DQ2/8 carriers and non-carriers without autoimmune disease, we hypothesize a potential genotype–microbiome–host axis. As key antigen-presenting molecules, HLA-DQ heterodimers display distinct peptide-binding preferences depending on genotype, thereby shaping T-cell responses ^[21–23]^. Many microbial peptides, particularly those derived from cell-wall and membrane proteins, are processed through this pathway. Accordingly, DQ haplotypes could impose divergent immune pressures, modulating strain-level microbial divergence in the gut and saliva. This divergence, in turn, may contribute to shifts in microbial functions and host metabolic outcomes. A concrete example is the pantothenate axis. Because humans cannot synthesize pantothenate, circulating levels are determined by dietary intake as well as microbial biosynthesis and consumption ^[19,24,25]^. In our cohort, serum pantothenate was significantly lower in DQ2/8 carriers. Similar nutrient intake between groups suggests that diet did not drive these differences, despite lacking direct pantothenate intake data. Metagenomic profiling revealed a coordinated enrichment of the entire coenzyme A biosynthetic apparatus in DQ2/8+ carriers. This shift was characterized by consistent increases in both the de novo pantothenate biosynthesis segments and the downstream CoA conversion pathways, collectively manifesting as a significantly elevated superpathway of coenzyme A biosynthesis I. This pattern suggests that the DQ2/8-associated microbial community, especially *B. obeum*, may act as a nutrient sink, actively funneling and retaining pantothenate intracellularly, leaving less net pantothenate available to the host. Given the central role of the pantothenate–CoA pathway in lipid metabolism ^[20]^, reduced pantothenate availability may contribute to the lower HDL-C observed in carriers, as clinical and experimental studies showed that pantethine supplementation improves lipid profiles ^[26,27]^. Consistently, we found a positive correlation between serum pantothenate and HDL-C. These interactions are summarized in a schematic model that integrates HLA-DQ–associated microbial divergence, vitamin metabolism, and host lipid physiology (Figure 5).

**Figure 5.**
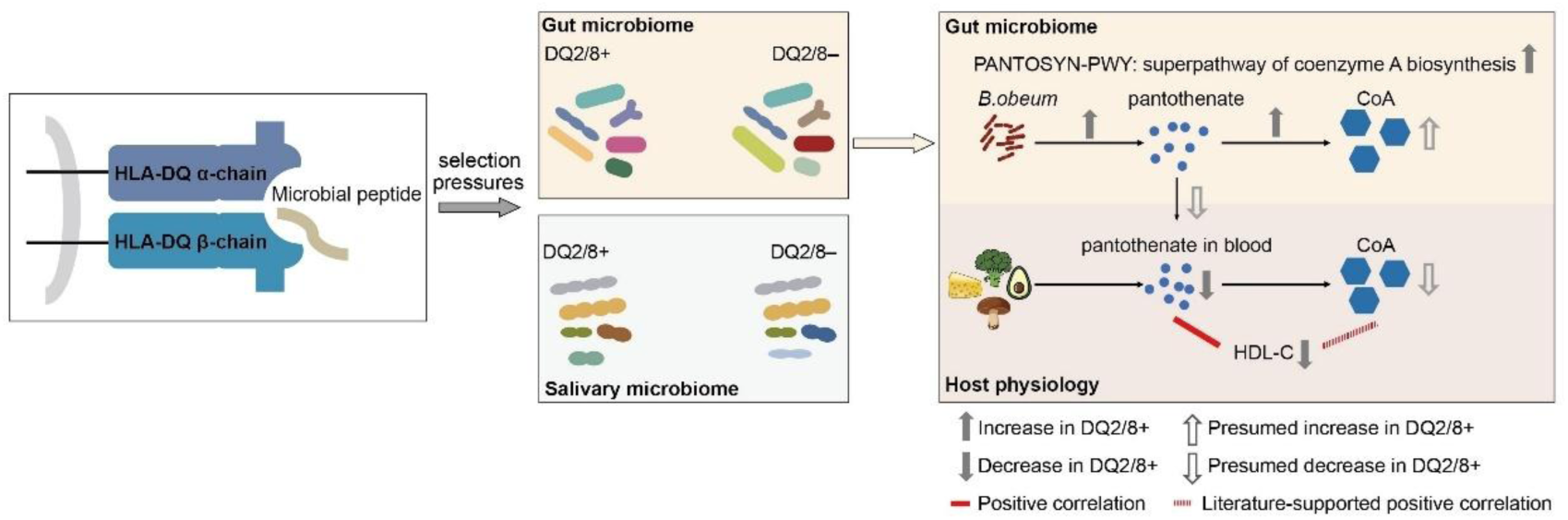
Schematic representation of the hypothesized HLA-DQ2/8–microbiome–host interaction axis. HLA-DQ heterodimers exhibit distinct peptide-binding preferences, imposing divergent immune selection pressures on colonizing microbes. This may result in strain-level divergence between DQ2/8 carriers (DQ2/8+) and non-carriers (DQ2/8–) in both gut and saliva microbiomes. In carriers, gut metagenomes show enrichment of superpathway of coenzyme A biosynthesis, phosphopantothenate biosynthesis and Coenzyme A biosynthesis, largely driven by *Blautia obeum*. These microbial adaptations may contribute to reduced host circulating pantothenate, diminished CoA availability, and lower HDL-C. Together, the model illustrates a potential genotype–microbiome–host axis linking HLA-DQ2/8-associated antigen presentation, strain-level microbial divergence, and host metabolic variation.

This study advances understanding of HLA-DQ–microbiome interactions in several ways. First, we benchmarked the effect size of HLA-DQ2/8 genotype against major host/environmental covariates in healthy adults without autoimmune disease, revealing that the genotype exceeded sex and BMI, though remained smaller than age and a few dietary factors, especially fiber. Our results suggest that HLA genotype should be considered as a key covariate in microbiome-based risk models and biomarker development. Second, strain-resolved metagenomics showed that certain key taxa (*e.g.*, *B. obeum*) differed primarily in functional genome content rather than species-level abundance, indicating intra-species adaptation to DQ-dependent immune niches. Third, by integrating metabolomic and clinical data, we propose a potential mechanistic chain linking HLA-DQ2/8-associated microbial functions (*e.g.*, CoA metabolism) to host phenotypes (*e.g.*, HDL-C), thereby connecting this genotype to systemic physiology. Finally, our adult cohort suggests that HLA-DQ-associated microbiome differences persist beyond early life, complementing previous infant and child studies ^[3,4,11–13]^.

Consistent with prior work in Western infants and children, our study in Asian adults without autoimmune disease also identified shared microbial features of HLA-DQ2/8 carriers. The features include a higher abundance of the butyrate producer *Faecalibacterium* and *C. catus*, as well as an enriched genomic potential for B-family vitamin and cofactor biosynthesis, specifically within the thiamine (B1) and pantothenate (B5)-coenzyme A pathways ^[13]^. However, important differences were evident. In schoolchildren, HLA-DQ2/8 risk haplotypes were associated with reduced Faith’s phylogenetic diversity ^[12]^, suggesting lower stability and resilience, whereas adult carriers here exhibited increased Shannon diversity. Berryman *et al*. reported that infants carrying risk haplotypes harbored gut microbiota enriched in pathogen-associated molecular patterns (PAMPs) and virulence genes, indicative of greater pro-inflammatory potential ^[13]^. In contrast, adult carriers here displayed fewer virulence factors. Moreover, Berryman *et al*. found a microbial gene signature centering on Gram-negative flagellar and O-antigen biosynthesis pathways ^[13]^, whereas the intra-species divergence in our cohort was predominantly localized in Gram-positive taxa, with microbial antigens primarily originating from peptidoglycan and membrane components. These discrepancies may reflect shifted ecological dynamics across developmental stages or population-specific backgrounds. Future studies utilizing longitudinal, multi-ethnic cohorts spanning from infancy to adulthood will be essential to disentangle these complex host–microbial interplays.

Building upon these observations, adult DQ2/8+ carriers here exhibited a seemingly healthier microbial profile, characterized by elevated diversity, reduced virulence, and abundant butyrate producers. This finding seems paradoxical given the well-established role of HLA-DQ2/8 as an autoimmune-predisposing factor. We speculate that this state may not reflect a true baseline of health, but rather a homeostatic high-risk set point ^[28,29]^. Supporting this hypothesis is the significantly elevated investment in the microbial pantothenate-CoA axis observed in carriers. This may sustain protective functions, such as butyrate synthesis via the butyryl-CoA:acetate CoA-transferase pathway, while simultaneously depleting systemic host pantothenate. Given the essential role of B-family vitamins in the regulation of host immunity, particularly in maintaining mucosal integrity and T-cell homeostasis ^[25,30]^, such a host-side deficiency could theoretically compromise immune-metabolic reserves and heighten autoimmune susceptibility. We hypothesize that while this compensatory adaptation buffers genetic risk, it maintains the system in a precarious, primed equilibrium. Consequently, a microbial buffer collapse ^[31–33]^ might occur under cumulative environmental stressors, potentially precipitating a transition toward autoimmunity. Ultimately, while our data suggest an interaction between HLA-DQ2/8 genotype, the microbiome, and host immune-metabolic regulation, the precise mechanisms linking genotype to disease susceptibility remain to be fully elucidated.

Beyond the gut microbiome, our results suggest that the influence of the HLA-DQ2/8 genotype is also reflected in the oral microbiome. Although the signals in the oral cavity were less pronounced than those in the gut, strain-level analysis nonetheless revealed significant divergence between genotypes. Given the established links between oral dysbiosis and systemic conditions such as cardiovascular disease, diabetes and rheumatoid arthritis ^[34–38]^, the consequences of HLA-driven selection at this secondary colonization site warrant further investigation. Furthermore, while this study focused on DQ2/8, exploring how other HLA haplotypes affect microbiomes remains a crucial open question. Addressing these gaps requires larger, multi-ethnic cohorts, combined multi-omic profiling, and mechanistic experimental validation. Such efforts will not only broaden our understanding of HLA–microbiome interactions, but also pave the way for personalized microbiome-based strategies to prevent and manage HLA-linked disease.

Several limitations of this study should be acknowledged. First, the modest sample size (N=60) and relatively small effect sizes mean that some biologically significant signals may have remained below the threshold of detection. Second, while the recruitment of a homogeneous cohort of Chinese university students effectively minimized environmental confounding, it potentially limits the generalizability of our findings to broader populations. Third, although our structural modeling using AlphaFold3 and NetMHCIIpan provides a compelling *in silico* framework for HLA–microbial peptide interactions, these predictions require future functional validation through *in vitro* binding assays and *in vivo* T-cell activation studies. Additionally, the mechanistic chain linking microbial genomic abundance to serum metabolites and lipid phenotypes involves several unverified steps that warrant further experimental validation. Fourth, our primary analysis grouped DQ2.2, DQ2.5, and DQ8 carriers together. This strategy increased statistical power and allowed us to identify microbial and metabolic features shared by DQ2/8 carriage, but it may have obscured haplotype-specific effects related to distinct peptide-binding repertoires and autoimmune-risk architectures. Larger allele-resolved cohorts will be required to distinguish shared DQ2/8-associated signatures from DQ2.2-, DQ2.5-, or DQ8-specific microbial patterns.

In summary, this study highlights that HLA-DQ2/8-related genetic risk may extend beyond a static host predisposition and be reflected in a dynamic microbiome–metabolic landscape, offering new insights into the interactions between the human immune system and its resident microbiota.

## Methods

### Study population and design

This investigation was nested within a single-arm, fixed-sequence (ABA) dietary intervention assessing the effects of gluten intake on microbiome and host physiology (registered with the Chinese Clinical Trial Registry, ChiCTR2100045383; registered on 14 April 2021). All participants completed two weeks of low-gluten intake, four weeks of high-gluten intake, and two weeks of low-gluten intake, in that specific order. Longitudinal sampling was conducted at two-week intervals. The present observational study is a genotype-stratified secondary analysis of this longitudinal cohort. It examines associations between HLA-DQ2/8 carrier status and host–microbiome features.

Participants were healthy undergraduate and graduate students recruited at Jiangnan University in 2021. Inclusion criteria included: age >18 years, regular dietary habit of rice as the staple food, no gluten intolerance, no history of chronic, metabolic, autoimmune, or gastrointestinal disease, no history of seizure symptoms or psychiatric disorders, no acute illness within 3 months, no participation in any special diet regimen or significant dietary changes in the past month, no use of antibiotics within the past 3 months, non-smokers, and not pregnant or lactating. All health-related conditions were self-reported by the participants. Participants were instructed to restrict habitual gluten intake to < 2 g/day. During the high-gluten phase, participants received a daily supplement of 17.35 g gluten. During low-gluten phases, iso-caloric soy-protein supplements were provided. A total of 60 individuals were included in the present analysis (see flow diagram in Figure S1A).

### Sample collection and clinical phenotyping

For each participant, five dietary records were collected, along with five stool and five saliva samples for microbiome profiling; three physical examinations were conducted, providing three serum samples for metabolomic profiling and three clinical phenotyping datasets (Figure S1B). Additionally, intestinal permeability testing was performed three times for a subgroup of 24 participants. All samples were transported immediately to the laboratory, aliquoted and stored at −80 °C until analysis. Comprehensive details are provided in the Supplementary Methods.

### Dietary assessment

Dietary records were obtained using a dedicated online management platform (WeChat Mini Program Banji Xiaoguanjia). Before and after each eating occasion, participants uploaded photographs of all meals, beverages, and snacks, accompanied by textual descriptions of dishes, portion sizes, and additional notes if applicable. Daily nutrient intakes including energy, macronutrients (carbohydrates, protein, fat, and fiber), micronutrients (vitamins A, C, E, B1, B2, B3, sodium, calcium, and iron), and cholesterol were estimated from the dietary records using the Chinese Food Composition Table (6th edition) ^[39]^. Participants recorded their diet over four consecutive days every two weeks, and the mean intake across these four days was used in subsequent analyses. Comprehensive details are provided in the Supplementary Methods.

### HLA-DQ genotyping and tTG IgA measurement

Total DNA was extracted from saliva using the DNeasy PowerSoil Pro Kit (Qiagen). Genotyping was performed with the SALSA MLPA Probemix P438-D2 HLA kit (MRC Holland). MLPA products were separated by capillary electrophoresis on an ABI 3730XL Genetic Analyzer (Applied Biosystems). Peak calling and dosage analysis were performed in Coffalyser.Net (MRC Holland). Haplotypes identified included HLA-DQ2.2, HLA-DQ2.5, HLA-DQ8, and HLA-DQ7.5. In individuals without these haplotypes, HLA-DQ6 and HLA-DQ9 were selected as representative haplotypes, given their high prevalence in Southern China (China Zhejiang Han pop 2, Allele Frequency Net Database ^[40]^).

Serum tissue transglutaminase (tTG) IgA was measured using a commercial anti-tTG ELISA kit (EUROIMMUN).

### Metagenomic sequencing and data processing

Host DNA in saliva was depleted by osmotic lysis of host cells followed by nuclease digestion of cell-free DNA, as previously described ^[41]^. DNA was then extracted from stool and host-depleted saliva samples using the DNeasy PowerSoil Pro Kit. Sequencing libraries were constructed with the QIAseq FX DNA Library Kit (Qiagen), yielding an average insert size of ∼350 bp. Circularized libraries were sequenced on the DNBSEQ-T7 platform (BGI) with 150 bp paired-end reads.

Raw sequence reads were processed with KneadData v0.6.1 ^[42]^ using default parameters to remove low-quality bases and host-derived reads. For stool samples, a median of 69.0 million clean reads was obtained (interquartile range [IQR] 66.8–72.5 million), and for saliva samples, 22.6 million (IQR 16.6–26.5 million). Taxonomic profiling was performed using MetaPhlAn v4.0 ^[43]^, and functional profiling was conducted with HUMAnN v3.0 ^[44]^. UniRef90 gene families were annotated to obtain Enzyme Commission (EC) numbers and Kyoto Encyclopedia of Genes and Genomes Orthology (KO). Virulence factors were annotated using the Virulence Factor Database (VFDB) ^[45]^.

### Strain-level phylogenetic and functional analysis

Intra-species phylogenetic divergence was assessed using anpan_pglmm from the R package *anpan* v0.3.0 ^[16]^, adjusting for potential confounders including age, sex, BMI, and dietary nutrient intake. Functional divergence was evaluated using anpan_pwy_ranef. All analyses were performed using default parameters. Comprehensive details are provided in the Supplementary Methods.

To identify microbial genes associated with HLA-DQ genotypes, we analyzed longitudinal gene presence–absence patterns across the five time points. A gene was considered stably present in an individual if detected in at least three time points. Gene prevalence was then calculated separately for the DQ2/8+ and DQ2/8– groups. Genes with a prevalence difference greater than 15% between groups were defined as differential, and subjected to Gene Ontology (GO) enrichment analysis using the *topGO* R package v2.52.0 ^[46]^. The background gene set for enrichment analysis included all genes of the tested species, which were obtained from the UniRef90 database. GO annotations were derived from UniProt-GOA annotations via UniProtKB (release 2025_02). Fisher’s exact test was used to determine statistical significance for GO term enrichment.

### HLA-DQ binding predictions and structural modeling

The microbial peptide binding affinity to HLA class II molecules was predicted using NetMHCIIpan v4.3 ^[47]^. Amino acid sequences for genes associated with HLA-DQ genotypes were retrieved from the UniProt database, fragmented into 15-mer peptides, and analyzed for binding affinity against each HLA-DQ molecule.

For structural predictions of the HLA–peptide complexes, AlphaFold3 was employed in complex-prediction mode ^[18]^. The α- and β-chain protein sequences of the HLA-DQ heterodimer were retrieved from the IPD-IMGT/HLA database ^[48]^. All predictions were carried out using default parameters. The quality of the generated models was evaluated using the pLDDT (predicted Local Distance Difference Test) score.

### Metabolomics

Metabolites were extracted from 100 µL of serum, and untargeted metabolomics were performed by liquid chromatography–mass spectrometry (LC–MS) (Supplementary Methods). A total of 21,011 MS features were initially extracted with Compound Discoverer v3.3.1.111. Missing chromatographic peaks were filled using the “Fill Gaps” node with default settings. Features were retained as metabolites if they were absent from blanks, exhibited an absolute mass deviation less than 5 ppm, and had a full match to at least one reference database (MzCloud, ChemSpider, or MassList). This procedure yielded 500 candidate metabolites. After manual curation utilizing the Human Metabolome Database (HMDB) ^[49]^ to exclude compounds irrelevant to human metabolism, 420 metabolites remained for downstream analysis. Peak areas were log 10-transformed for downstream analyses.

Stool short-chain fatty acids (SCFAs) were quantified using gas chromatography–mass spectrometry (GC–MS) as previously described ^[50]^.

### Statistical analysis

All statistical analyses were performed in R v4.3.2.

To generate a stable representative profile for each individual, all microbiome, dietary, metabolomic, phenotypic, and questionnaire-derived variables were averaged across the five sampling time points for each participant unless otherwise specified. This aggregation resulted in a single, consolidated observation per participant for downstream cross-sectional analysis.

Global differences in gut microbiome profiles (species, pathways, KOs, GOs, ECs) were assessed using permutational multivariate analysis of variance (PERMANOVA), implemented in the adonis2 function of the *vegan* R package v2.6-4 with the option *by="margin"*.

Microbial taxonomic and functional relative abundances were transformed using the arcsine square-root transformation prior to linear modeling. Differential species, metabolites, and host phenotypes between HLA-DQ2/8 carriers and non-carriers were identified using multiple linear regression models with the lm function in R. Covariates included age, sex, and BMI in all models. For microbiome analyses, dietary nutrient intake variables were additionally included to account for potential dietary confounding. These variables were selected based on an evaluation of multicollinearity using the variance inflation factor (VIF) from the *car* R package v3.1.5 ^[51]^. Specifically, fat, carbohydrates, fiber, vitamin A, vitamin B3, vitamin C, cholesterol, and calcium were included in the final models, as all exhibited VIF values < 5.

Multiple testing correction was performed using the Benjamini–Hochberg method, controlling the false discovery rate (FDR) at 0.1 unless otherwise specified (reported as adjusted values).

A previous version of this manuscript has been deposited as a preprint in bioRxiv ^[52]^.

## Acknowledgements

We thank all participants for their contribution to this study. We are also grateful to Professor Yanyi Huang for support with metabolomics.

## Contributors

Chuanwen Kan: Formal analysis, Investigation, Visualization, Writing – original draft.

Mo Hu: Methodology, Investigation.

Ning Wang: Data curation, Investigation.

Qianhui Zhang: Data curation.

Lingxiao Chaihu: Investigation.

Xuan Jiang: Investigation.

Chun Wang: Investigation.

Wenwei Lu: Resources.

Guanbo Wang: Supervision.

Mingkun Li: Supervision.

Li Zhang: Conceptualization, Investigation, Data curation, Writing – review & editing, Project administration, Funding acquisition.

## Funding

This work was supported by the Youth Innovation Promotion Association of Chinese Academy of Sciences (grant no. 2021097 to L.Z.). The funder had no role in study design, data collection, data analysis, data interpretation, writing of the manuscript, or the decision to submit the work for publication.

## Competing interests

None declared

## Patient consent for publication

Not required

## Patient and Public Involvement

Patients or the public were not involved in the design, conduct, reporting, or dissemination of this research.

## Ethics approval

The study was conducted in accordance with the Declaration of Helsinki. The study was approved by the Institutional Review Board at Jiangnan University Affiliated Hospital on May 8, 2021 (Ethics Approval No.: LS2021025). Written informed consent was obtained from all participants.

## Data availability statement

All materials used in this study are original. Sequencing data were deposited in the Genome Sequence Archive at the China National Center for Bioinformation, under accession CRA029700. The preview link is https://ngdc.cncb.ac.cn/gsa/s/eqLKhSDt. Scripts used for the analyses are available at https://github.com/chwenkan/HLA-DQ28-genotype/tree/main.

## Declaration of generative AI use

During the preparation of this work, the authors used ChatGPT to assist with language editing. After using this tool, the authors reviewed and edited the content as needed and take full responsibility for the content of the published article.

## Notes

https://github.com/chwenkan/HLA-DQ28-genotype/tree/main

### Competing Interest Statement

The authors have declared no competing interest.

### Summary of Updates

This revised version updates the title, abstract, main text, methods, results, discussion, figures, tables, and supplemental materials. The analysis strategy was refined, cohort characterization and robustness analyses were added, and the interpretation was revised to more cautiously describe the association between HLA-DQ2/8 genotype, strain-level microbiome divergence, serum pantothenate, and HDL-C.

