## Supplementary material for "HLA-DQ2/8 genotype is associated with strain-level gut microbiome divergence and lower serum pantothenate in healthy adults"

**Content:**

Supplementary Methods

Supplementary Table 1

Supplementary Table 2

Supplementary Table 3

Supplementary Table 4

Supplementary Figure 1

Supplementary Figure 2

Supplementary Figure 3

**Supplementary Methods**

**Sample collection and clinical phenotyping**

Participants were instructed in standardized procedures for stool and saliva collection using sterile containers. Stool consistency was self-evaluated using the Bristol Stool Form Scale. Saliva samples were obtained in the morning, at least 30 minutes after eating or drinking. All samples were transported immediately to the laboratory, aliquoted and stored at −80 °C until analysis. Stool pH was measured using a calibrated pH meter.

On the evening prior to physical examinations, participants were asked to refrain from alcohol, coffee, tea, and dietary supplements (*e.g.*, vitamins, fish oil) other than those supplied in the trial, and to avoid strenuous physical activity. All examinations were performed in the fasting state the following morning. Standard clinical assessments included complete blood count, serum biochemistry, urinalysis, blood pressure, and heart rate. Anthropometric measures including height, weight, body fat, and muscle mass were recorded using calibrated instruments.

For intestinal permeability testing, participants fasted overnight, emptied their bladders, and then ingested a solution containing 10 g lactulose and 4 g mannitol dissolved in 200 mL water. Urine was collected over the subsequent 4 h, centrifuged at 4,500 g for 10 min, and 300 μL of the supernatant was filtered through a 0.22 μm aqueous membrane. Lactulose and mannitol concentrations were quantified by high-performance anion-exchange chromatography with pulsed amperometric detection (HPAE-PAD; ICS-5000+SP-5, Thermo Fisher Scientific), and the lactulose/mannitol ratio was calculated as a marker of intestinal mucosal permeability.

**Dietary assessment**

All dietary records were curated by two trained researchers, who standardized the entries into structured items including dish/food name, source (university dining halls, off-campus restaurants, packaged foods, raw fruits and vegetables, or study-supplied supplements), and standardized portion size. All entries were independently reviewed and double-checked by a registered dietitian.

Nutrient intake was estimated using a two-step approach. For freshly prepared dishes and foods, names were converted into standardized ingredient lists and corresponding ingredient quantities using an in-house recipe-based ingredient composition database, and nutrient values were subsequently derived from these ingredients using the Chinese Food Composition Table (6th edition). For packaged foods, nutrient content was calculated directly from the manufacturer-provided ingredient and nutrition labels. This procedure provided daily estimates of energy, macronutrients (carbohydrates, protein, fat, and fiber), micronutrients (vitamins A, C, E, B1, B2, B3, sodium, calcium, and iron), and cholesterol. Ingredients were further classified into 18 food groups and 98 subgroups based on the food categories in the Chinese Food Composition Table, and the number of food groups/subgroups consumed was used as indicators of dietary diversity.

**Strain-level phylogenetic and functional analysis**

Intra-species phylogenetic divergence was assessed using anpan_pglmm from the R package *anpan* v0.3.0, following the modeling framework described previously. Briefly, species-specific gene presence–absence matrices derived from HUMAnN3 gene-family profiling were used to construct strain-level phylogenetic trees. A phylogenetic generalized linear mixed model (pGLMM) was then fitted to estimate intra-species phylogenetic divergence while accounting for host covariates and phylogenetically structured variation. Species relative abundance across samples was included as an additional fixed-effect covariate to account for abundance-related effects, alongside age, sex, BMI, and dietary variables.

Functional divergence was evaluated using anpan_pwy_ranef with default parameters. This approach applies a pathway random-effects model to identify pathways whose abundances differ between groups while accounting for the correlation between pathway abundance and species abundance. Prior to modeling, pathway abundances were log-transformed and zero values were excluded. The relationship between species abundance and pathway abundance was estimated across all pathways within a species. Pathway-specific intercepts and slopes were modeled as random effects, allowing partial pooling across pathways.

**Metabolomics**

Metabolites were extracted from 100 µL of serum with 400 µL of cold methanol. Precipitated proteins were removed by centrifugation at 14,000 g for 15 min at 4 °C, and 200 µL of the supernatant was collected and dried in a vacuum concentrator. The dried extracts were reconstituted in 200 µL of 0.1% formic acid in water prior to analysis. Equal 20 µL aliquots of each sample were pooled to generate a quality control (QC) sample.

Liquid chromatography–mass spectrometry (LC–MS) was performed on a Vanquish Neo microflow LC system (Thermo Fisher Scientific) coupled to an Orbitrap Exploris 480 mass spectrometer (Thermo Fisher Scientific) equipped with an electrospray ionization source. Metabolites were separated on a 1.0 mm × 150 mm reversed-phase column at a flow rate of 50 µL/min using the following linear gradient: 0–4 min, 0% B; 4–17 min, 0–100% B; 17–20 min, 100% B. Mobile phase A was 0.1% formic acid in water, and mobile phase B was 0.1% formic acid in acetonitrile. The mass spectrometer was first operated in data-dependent acquisition mode (m/z 70–1050; MS resolution 60,000; MS/MS resolution 30,000) with AcquireX control using the QC sample to generate a comprehensive spectral library. Subsequently, randomized study samples were analyzed in MS1-only mode (resolution 120,000) for quantification.

**Supplementary Table 1. Comprehensive Physiological, Biochemical, and Psychosocial Characteristics for Cohort Homogeneity Assessment.**

| **Class** | **Feature** | **DQ2/8– (n=32)** | **DQ2/8+ (n=28)** | **P value*** | **p.adj** |
| --- | --- | --- | --- | --- | --- |
| Body composition | Skeletal muscle mass | 20.8 [19.9-21.9] | 19.7 [18.9-20.8] | 0.055 | 0.672 |
| Body composition | Body fat percentage | 26.8 [21.6-29.7] | 24.9 [22.1-28.5] | 0.456 | 0.964 |
| Renal function | Creatinine | 54.9 [52.0-57.2] | 55.5 [52.0-61.0] | 0.793 | 0.964 |
| Renal function | Blood urea nitrogen | 3.8 [3.2-4.4] | 4.2 [3.5-4.7] | 0.109 | 0.845 |
| Liver function | Total bilirubin | 14.8 [12.4-17.8] | 13.6 [11.6-17.8] | 0.417 | 0.964 |
| Liver function | Direct bilirubin | 4.8 [4.2-5.5] | 4.4 [3.5-5.8] | 0.482 | 0.964 |
| Liver function | Alanine aminotransferase | 11.6 [9.7-14.6] | 11.2 [9.0-14.2] | 0.924 | 0.964 |
| Liver function | Aspartate aminotransferase | 18.9 [16.8-19.8] | 17.8 [16.6-19.4] | 0.238 | 0.964 |
| Liver function | Alkaline phosphatase | 46.0 [42.5-50.0] | 44.8 [40.3-50.7] | 0.804 | 0.964 |
| Liver function | Gamma-glutamyl transferase | 13.3 [10.9-15.9] | 13.6 [11.0-15.1] | 0.931 | 0.964 |
| Serum proteins | Albumin | 46.3 [45.3-47.4] | 46.4 [45.0-47.4] | 0.587 | 0.964 |
| Serum proteins | Total protein | 72.5 [71.2-75.2] | 73.0 [71.7-74.0] | 0.915 | 0.964 |
| Red blood cell | Red blood cell count | 4.4 [4.2-4.6] | 4.3 [4.1-4.5] | 0.185 | 0.964 |
| Red blood cell | Hemoglobin concentration | 130.7 [127.0-135.3] | 129.0 [123.3-133.2] | 0.243 | 0.964 |
| Red blood cell | Hematocrit | 40.4 [39.5-41.6] | 40.5 [37.9-41.2] | 0.458 | 0.964 |
| Red blood cell | Mean corpuscular hemoglobin concentration | 325.7 [318.3-328.8] | 320.3 [317.8-328.2] | 0.477 | 0.964 |
| Red blood cell | Mean corpuscular hemoglobin | 29.6 [28.5-30.3] | 29.2 [28.8-30.4] | 0.791 | 0.964 |
| Red blood cell | Mean corpuscular volume | 91.3 [88.4-94.9] | 91.5 [88.9-93.8] | 0.819 | 0.964 |
| Red blood cell | Red cell distribution width–standard deviation | 40.5 [39.3-41.1] | 40.2 [38.8-41.9] | 0.933 | 0.964 |
| Red blood cell | Red cell distribution width–coefficient of variation | 12.2 [11.9-12.7] | 12.3 [12.0-12.7] | 0.921 | 0.964 |
| Urinalysis | Urine specific gravity | 1.0 [1.0-1.0] | 1.0 [1.0-1.0] | 0.293 | 0.964 |
| Urinalysis | Uric acid | 319.7 [287.7-356.7] | 318.0 [275.7-366.0] | 0.659 | 0.964 |
| Urinalysis | Urine pH | 5.8 [5.5-6.2] | 5.8 [5.5-6.2] | 0.982 | 0.982 |
| Psychosocial and lifestyle traits | Sleep duration (questionnaire) | 7.0 [7.0-7.4] | 7.0 [7.0-7.4] | 0.766 | 0.964 |
| Psychosocial and lifestyle traits | Sleep quality (questionnaire) | 4.0 [3.4-4.2] | 4.0 [3.6-4.0] | 0.707 | 0.964 |
| Psychosocial and lifestyle traits | Attention level (questionnaire) | 3.0 [3.0-3.2] | 3.0 [3.0-3.2] | 0.771 | 0.964 |
| Psychosocial and lifestyle traits | Energy level (questionnaire) | 3.0 [3.0-3.1] | 3.2 [3.0-3.4] | 0.065 | 0.672 |
| Psychosocial and lifestyle traits | Negative emotions score (questionnaire) | 3.0 [3.0-3.4] | 3.2 [3.0-3.4] | 0.528 | 0.964 |
| Psychosocial and lifestyle traits | Mental health score (questionnaire) | 7.7 [7.2-8.4] | 8.2 [7.2-8.4] | 0.427 | 0.964 |
| Physiological health | Physical health score (questionnaire) | 7.7 [7.2-8.4] | 7.8 [7.4-8.4] | 0.476 | 0.964 |

*P values were calculated using the Wilcoxon rank-sum test. The data are presented as median [IQR].

**Supplementary Table 2. Metabolites theoretically linked to the differentially abundant microbial pathways.**

| **Metabolite** | **Detected in serum metabolomics** | **Associated microbial pathway(s)** | **Number of matched pathways** |
| --- | --- | --- | --- |
| Pantothenate | Yes | PANTO-PWY; PANTOSYN-PWY | 2 |
| Inosine | Yes | PWY-6609 | 1 |
| Hypoxanthine | Yes | PWY-6609 | 1 |
| L-histidine | Yes | HISTSYN-PWY; TRNA-CHARGING-PWY | 2 |
| L-phenylalanine | Yes | COMPLETE-ARO-PWY; TRNA-CHARGING-PWY | 2 |
| L-tyrosine | Yes | COMPLETE-ARO-PWY; TRNA-CHARGING-PWY | 2 |
| Tryptophan | Yes | COMPLETE-ARO-PWY | 1 |
| Anthranilic acid | Yes | COMPLETE-ARO-PWY | 1 |
| L-threonine | Yes | TRNA-CHARGING-PWY | 1 |
| L-glutamic acid | Yes | TRNA-CHARGING-PWY | 1 |
| Asparagine | Yes | TRNA-CHARGING-PWY | 1 |
| Valine | Yes | TRNA-CHARGING-PWY | 1 |
| L-methionine | Yes | TRNA-CHARGING-PWY | 1 |
| L-alanine | Yes | TRNA-CHARGING-PWY | 1 |
| Guanosine triphosphate | No | RIBOSYN2-PWY; PWY-6700 | 2 |
| Riboflavin | No | RIBOSYN2-PWY | 1 |
| Flavin mononucleotide (FMN) | No | RIBOSYN2-PWY | 1 |
| Flavin adenine dinucleotide (FAD) | No | RIBOSYN2-PWY | 1 |
| Pantoate | No | PANTO-PWY; PANTOSYN-PWY | 2 |
| β-Alanine | No | PANTO-PWY; PANTOSYN-PWY | 2 |
| 4′-Phosphopantothenate | No | COA-PWY; PANTOSYN-PWY | 2 |
| Pantothenoylcysteine | No | COA-PWY; PANTOSYN-PWY | 2 |
| 4′-Phosphopantothenoylcysteine | No | COA-PWY; PANTOSYN-PWY | 2 |
| Pantetheine | No | COA-PWY; PANTOSYN-PWY | 2 |
| 4′-Phosphopantetheine | No | COA-PWY; PANTOSYN-PWY | 2 |
| Coenzyme A | No | COA-PWY; PANTOSYN-PWY | 2 |
| 4-Amino-2-methyl-5-pyrimidinemethanol | No | PWY-6897 | 1 |
| Thiamine phosphate | No | PWY-6897 | 1 |
| Thiamine diphosphate | No | PWY-6897 | 1 |
| Bicarbonate | No | PWY-7790; PWY-7791; PWY-5686 | 3 |
| L-Glutamine | No | PWY-7790; PWY-7791; PWY-5686 | 3 |
| L-Aspartate | No | PWY-7790; PWY-7791; PWY-5686 | 3 |
| Carbamoyl phosphate | No | PWY-7790; PWY-7791; PWY-5686 | 3 |
| N-Carbamoyl-L-aspartate | No | PWY-7790; PWY-7791; PWY-5686 | 3 |
| Dihydroorotate | No | PWY-7790; PWY-7791; PWY-5686 | 3 |
| Orotidine 5′-monophosphate (OMP) | No | PWY-7790; PWY-7791; PWY-5686 | 3 |
| Uridine 5′-monophosphate (UMP) | No | PWY-7790; PWY-7791; PWY-5686 | 3 |
| Adenine | No | PWY-6609 | 1 |
| Adenosine | No | PWY-6609 | 1 |
| Adenosine monophosphate (AMP) | No | PWY-6609 | 1 |
| Inosine monophosphate (IMP) | No | PWY-6609 | 1 |
| 7-Cyano-7-deazaguanine (preQ0) | No | PWY-6700 | 1 |
| 7-Aminomethyl-7-deazaguanine (preQ1) | No | PWY-6700 | 1 |
| Epoxyqueuosine | No | PWY-6700 | 1 |
| Queuosine | No | PWY-6700 | 1 |
| Histidinol phosphate | No | HISTSYN-PWY | 1 |
| Histidinol | No | HISTSYN-PWY | 1 |
| Phosphoenolpyruvate | No | ARO-PWY; PWY-6163; COMPLETE-ARO-PWY | 3 |
| D-Erythrose 4-phosphate | No | ARO-PWY; COMPLETE-ARO-PWY | 2 |
| 3-Deoxy-D-arabino-heptulosonate 7-phosphate (DAHP) | No | ARO-PWY; COMPLETE-ARO-PWY | 2 |
| 3-Dehydroquinate | No | ARO-PWY; PWY-6163; COMPLETE-ARO-PWY | 3 |
| 3-Dehydroshikimate | No | ARO-PWY; PWY-6163; COMPLETE-ARO-PWY | 3 |
| Shikimate | No | ARO-PWY; PWY-6163; COMPLETE-ARO-PWY | 3 |
| Shikimate 3-phosphate | No | ARO-PWY; PWY-6163; COMPLETE-ARO-PWY | 3 |
| 5-Enolpyruvylshikimate 3-phosphate (EPSP) | No | ARO-PWY; PWY-6163; COMPLETE-ARO-PWY | 3 |
| Chorismate | No | ARO-PWY; PWY-6163; COMPLETE-ARO-PWY | 3 |
| Prephenate | No | COMPLETE-ARO-PWY | 1 |
| Phenylpyruvate | No | COMPLETE-ARO-PWY | 1 |
| 4-Hydroxyphenylpyruvate | No | COMPLETE-ARO-PWY | 1 |

**Supplementary Table 3. Comparison of downstream phenotypic outcome features between HLA-DQ2/8+ and HLA-DQ2/8– individuals.**

| **Class** | **Feature** | **Mean** | **Mean** | **Coefficient** | **P* value** | **p.adj** |
| --- | --- | --- | --- | --- | --- | --- |
|  |  | **(DQ2/8–)** | **(DQ2/8+)** |  |  |  |
| Gut health | Bowel movement frequency | 0.924 | 0.827 | -0.563 | 0.072 | 0.446 |
| Gut health | Bristol stool scale | 3.66 | 4.048 | 0.304 | 0.285 | 0.727 |
| Gut health | Gut permeability | 0.018 | 0.018 | -0.156 | 0.762 | 0.872 |
| Gut health | Stool PH | 6.423 | 6.378 | -0.236 | 0.397 | 0.770 |
| Fecal SCFA | Butyric acid | 47.134 | 63.846 | 0.544 | 0.058 | 0.446 |
| Fecal SCFA | Isovaleric acid | 6.925 | 7.921 | 0.474 | 0.099 | 0.512 |
| Fecal SCFA | Isobutyric acid | 5.642 | 6.632 | 0.391 | 0.194 | 0.668 |
| Fecal SCFA | Valeric acid | 6.158 | 7.448 | 0.323 | 0.260 | 0.727 |
| Fecal SCFA | Propionic acid | 92.680 | 111.297 | 0.298 | 0.305 | 0.727 |
| Fecal SCFA | Caproic acid | 2.130 | 2.412 | 0.244 | 0.430 | 0.770 |
| Fecal SCFA | Acetic acid | 242.489 | 251.728 | 0.121 | 0.674 | 0.872 |
| Cardiovascular | Heart rate | 80.406 | 75.304 | -0.658 | 0.015 | 0.446 |
| Cardiovascular | Diastolic blood pressure | 70.649 | 69.229 | -0.205 | 0.447 | 0.770 |
| Cardiovascular | Systolic blood pressure | 110.67 | 110.554 | -0.193 | 0.395 | 0.770 |
| Blood glucose | Blood glucose | 4.849 | 4.955 | 0.146 | 0.569 | 0.872 |
| Immune cell outcomes | White blood cell count | 5.653 | 5.336 | -0.079 | 0.776 | 0.872 |
| Immune cell outcomes | Lymphocyte count | 2.094 | 2.134 | 0.029 | 0.915 | 0.932 |
| Immune cell outcomes | Lymphocyte percentage | 38.008 | 40.793 | 0.489 | 0.060 | 0.446 |
| Immune cell outcomes | Monocyte count | 0.341 | 0.336 | -0.388 | 0.144 | 0.574 |
| Immune cell outcomes | Monocyte percentage | 6.146 | 6.254 | -0.111 | 0.680 | 0.872 |
| Immune cell outcomes | Neutrophil count | 3.088 | 2.748 | -0.218 | 0.400 | 0.770 |
| Immune cell outcomes | Neutrophil percentage | 53.519 | 50.808 | -0.378 | 0.148 | 0.574 |
| Immune cell outcomes | Basophil count | 0.023 | 0.019 | -0.524 | 0.055 | 0.446 |
| Immune cell outcomes | Basophil percentage | 0.416 | 0.351 | -0.024 | 0.932 | 0.932 |
| Immune cell outcomes | Eosinophil count | 0.107 | 0.1 | 0.138 | 0.627 | 0.872 |
| Immune cell outcomes | Eosinophil percentage | 1.923 | 1.81 | 0.296 | 0.301 | 0.727 |
| Platelet | Plateletcrit | 0.252 | 0.247 | -0.090 | 0.748 | 0.872 |
| Platelet | Mean platelet volume | 10.851 | 10.725 | -0.071 | 0.788 | 0.872 |
| Platelet | Platelet count | 230.322 | 232.723 | -0.093 | 0.736 | 0.872 |
| Platelet | Platelet large cell ratio | 30.894 | 30.135 | -0.032 | 0.903 | 0.932 |
| Platelet | Platelet distribution width | 12.381 | 12.265 | 0.089 | 0.734 | 0.872 |

*P values were derived from multiple linear regression models adjusted for age, sex, BMI. SCFA, short-chain fatty acid.

**Supplementary Table 4. Evaluation of genotype-by-diet interactions for key microbial and host features using linear mixed-effects models.**

| **Class** | **Feature** | **P_interaction_*** | **p.adj** |
| --- | --- | --- | --- |
| Gut microbiome | Shannon index | 0.068 | 0.585 |
| Gut microbiome | Virulence genes | 0.240 | 0.585 |
| Gut microbiome | *Coprococcus catus* | 0.135 | 0.585 |
| Gut microbiome | *Alistipes shahii* | 0.849 | 0.849 |
| Gut microbiome | *Alistipes putredinis* | 0.751 | 0.778 |
| Gut microbiome | *Lachnospiraceae bacterium* | 0.377 | 0.585 |
| Gut microbiome | *Roseburia faecis* | 0.539 | 0.645 |
| Gut microbiome | *Klebsiella pneumoniae* | 0.206 | 0.585 |
| Gut microbiome | *Roseburia hominis* | 0.356 | 0.585 |
| Gut microbiome | *Gordonibacter pamelaeae* | 0.290 | 0.585 |
| Gut microbiome | *Oscillibacter* sp ER4 | 0.352 | 0.585 |
| Gut microbiome | PWY-6163: chorismate biosynthesis from 3-dehydroquinate | 0.444 | 0.585 |
| Gut microbiome | TRNA-CHARGING-PWY: tRNA charging | 0.416 | 0.585 |
| Gut microbiome | ARO-PWY: chorismate biosynthesis I | 0.425 | 0.585 |
| Gut microbiome | COMPLETE-ARO-PWY: superpathway of aromatic amino acid biosynthesis | 0.426 | 0.585 |
| Gut microbiome | HISTSYN-PWY: L-histidine biosynthesis | 0.416 | 0.585 |
| Gut microbiome | PWY-6700: queuosine biosynthesis I (de novo) | 0.241 | 0.585 |
| Gut microbiome | PWY-6609: adenine and adenosine salvage III | 0.600 | 0.669 |
| Gut microbiome | PWY-7790: UMP biosynthesis II | 0.393 | 0.585 |
| Gut microbiome | PWY-7791: UMP biosynthesis III | 0.393 | 0.585 |
| Gut microbiome | PWY-5686: UMP biosynthesis I | 0.404 | 0.585 |
| Gut microbiome | PWY-6897: thiamine diphosphate salvage II | 0.556 | 0.645 |
| Gut microbiome | PANTOSYN-PWY: superpathway of coenzyme A biosynthesis I (bacteria) | 0.234 | 0.585 |
| Gut microbiome | COA-PWY: coenzyme A biosynthesis I (prokaryotic) | 0.502 | 0.633 |
| Gut microbiome | PANTO-PWY: phosphopantothenate biosynthesis I | 0.165 | 0.585 |
| Gut microbiome | RIBOSYN2-PWY: flavin biosynthesis I (bacteria and plants) | 0.642 | 0.690 |
| Serum metabolites | Pantothenate | 0.235 | 0.585 |
| Phenotype | High-density lipoprotein cholesterol | 0.259 | 0.585 |

*P values denote the significance of the genotype-by-diet interaction term. Statistical estimates were derived from linear mixed-effects models with participant ID as a random effect to account for the longitudinal, repeated-measures design. Fixed effects included HLA-DQ genotype, dietary intervention phase (baseline, low-gluten, and high-gluten time points), and their interaction, adjusting for age, sex, and BMI.

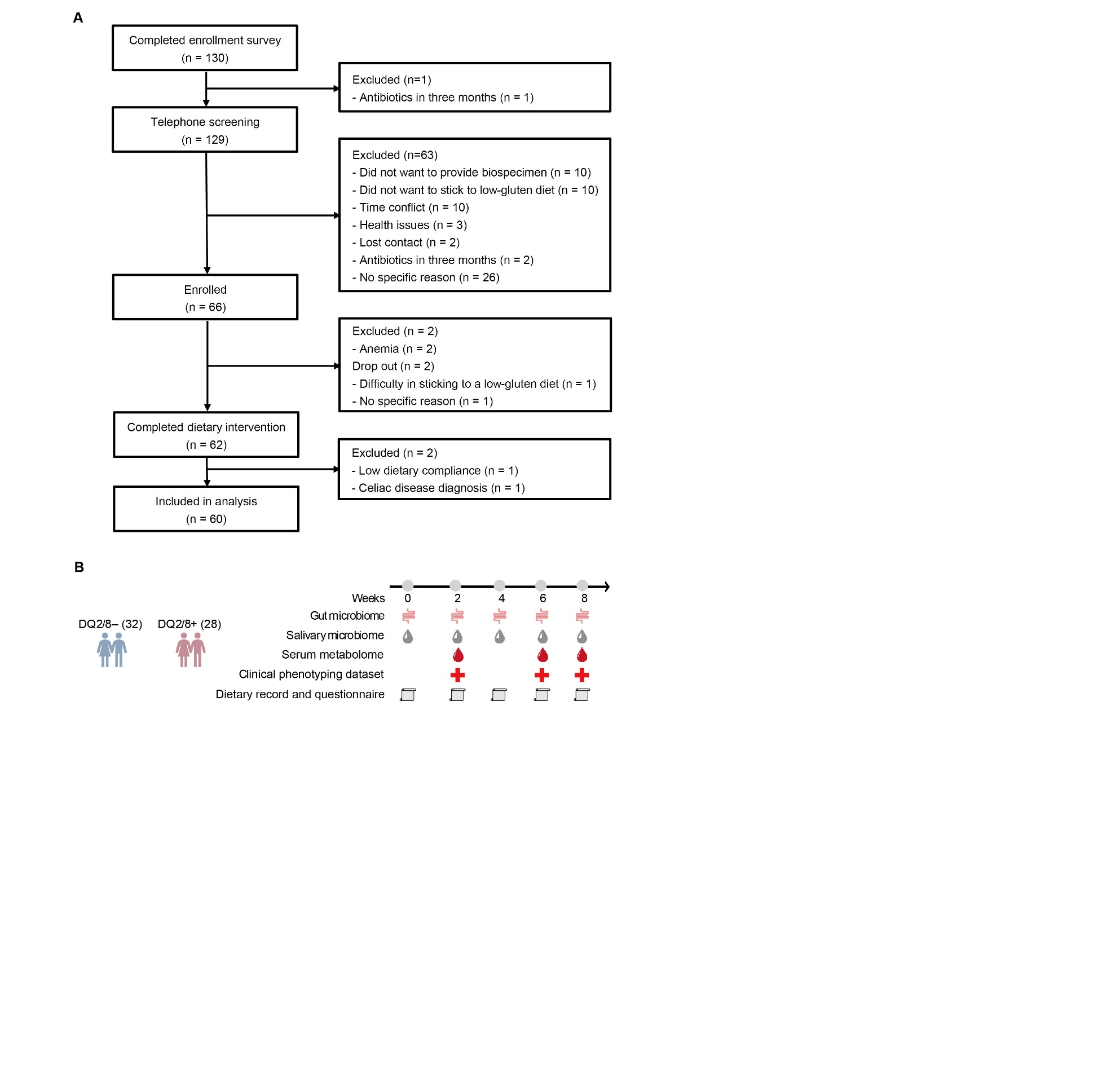

**Supplementary Figure 1. Study design and participant flow.**

(A) Flow diagram showing participant enrollment, exclusions, and completion. (B) Longitudinal sampling scheme and types of data collected.

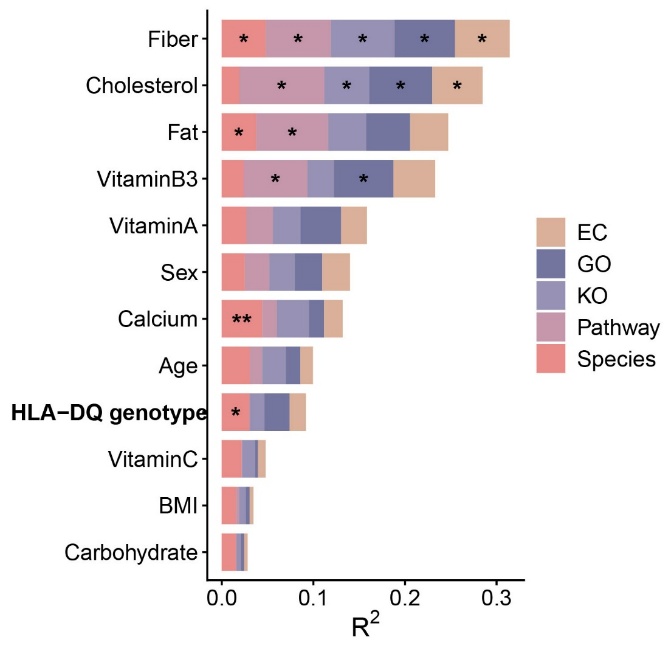

**Supplementary Figure 2. Comprehensive analysis of host factors associated with gut microbiome variation.**

All 12 analyzed host demographic and dietary covariates ranked by their effect size (Adonis R²) across five microbial profiles. Asterisks indicate statistical significance (*p<0.05; **p<0.01).

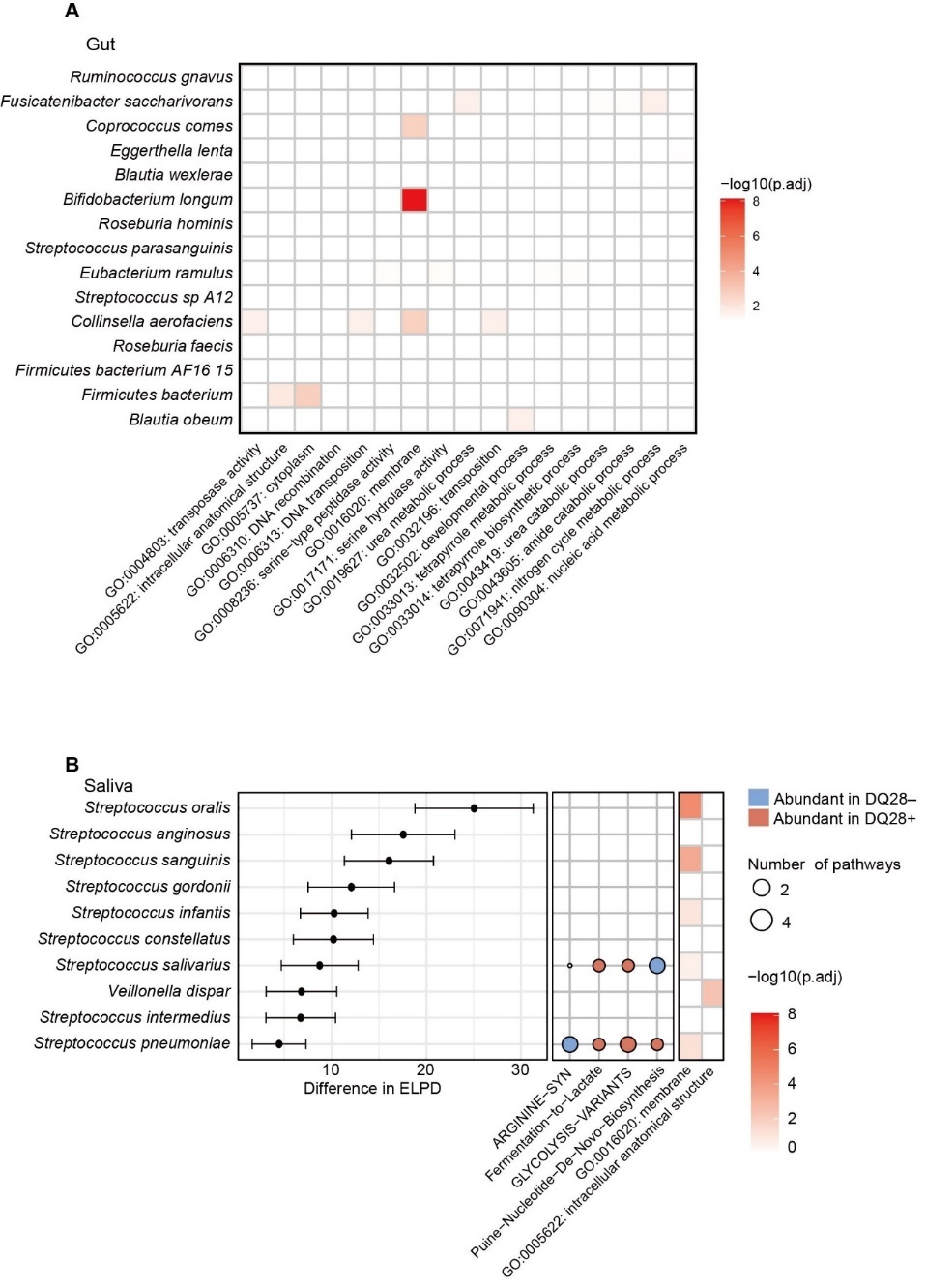

**Supplementary Figure 3. Extended Gene Ontology enrichment of gut microbial genes and intra-species divergence in the salivary microbiome**

(A) Heatmap of Gene Ontology (GO) enrichment of gut microbial genes showing differential prevalence between the DQ2/8+ and DQ2/8– groups within individual taxa. All significant GO terms are shown. (B) Salivary microbial intra-species divergence linked to DQ2/8 genotype. Left: Phylogenetic divergence between genotypes, as indicated by differences in expected log pointwise predictive density (ΔELPD) computed by anpan. Species with ΔELPD > 4 are shown. Center: Pathways showing genotype-associated divergences across species. Only pathways detected in more than five significant pathway-species associations are displayed. Right: Heatmap of Gene Ontology (GO) enrichment of salivary microbial genes showing differential prevalence between genotypes within individual taxa.
